# BAK knockdown delays bleaching and alleviates oxidative DNA damage in a reef-building coral

**DOI:** 10.1101/2024.03.14.585106

**Authors:** Eva Majerová, Camryn Steinle, Crawford Drury

## Abstract

Climate change is rapidly pushing coral reefs towards extinction. Efforts to restore and protect these ecosystems using resilient corals are increasing, but our understanding of cellular mechanisms of thermal resilience and trade-offs remains limited. Here, we demonstrate downregulation of pa-BAK slows bleaching and stabilizes the symbiosis during acute heat stress in *Pocillopora acuta*. Counterintuitively, oxidative DNA damage – a hallmark of the thermal stress response in corals – was prevented in corals with such “forced” symbiosis stability, possibly through alterations in mitochondrial ROS release. Using expression data of genes representing various stress-response pathways, we propose a model that coral heat stress response is mediated via the AMPK/Foxo3/Nrf2 signaling network. Developing our understanding of heat-stress defense mechanisms promoting stability in the coral-algal symbiosis is fundamental for effective modern coral reef restoration practices based on improving system resilience.

## Introduction

Ocean temperatures are virtually certain to increase, leading to more frequent and intense warming events that result in the breakdown of the coral-algal symbiosis (*1*, *2*). This trajectory compromises the structure and function of coral reefs and threatens their long-term persistence as the symbiotic relationship is the energetic foundation of reef-building corals (*3*). Despite years of research, the molecular and cellular mechanisms underpinning the initiation, maintenance, and disruption of this symbiosis are still largely unknown. Multiple mechanisms have been proposed (*4*), including programmed cell death pathways (PCD; apoptosis and symbiophagy), which play an important role in natural conditions and laboratory-induced bleaching experiments (*5–11*). These pathways involve the interaction of “pro-life” and “pro-death” proteins that inhibit or activate the execution of the pathway through posttranslational modifications and physical protein-protein interactions (*12*). BAK and BAX are two major apoptosis proteins that heterodimerize upon pro-death signaling, causing immediate permeabilization of the mitochondrial membrane, release of cytochrome c, fragmentation of mitochondria, and activation of caspases – proteases that degrade the content of the cell (*13*). Each is also involved in the autophagy cascade; both pathways are closely interconnected and can compensate for each other in certain conditions (*8*, *13*, *14*). In corals pre-exposed to sublethal temperature stress, the expression of the pro-survival gene pa-*Bcl-2* increases relative to pro-death genes pa-*Bak* and pa-*Bax* during the first 24h hours of acute thermal stress, but the inhibition of pa-*Bcl-2* negates this acquired beneficial phenotype (*5*).

Since inhibition of pa-Bcl-2 increased the bleaching rate of more resilient corals, the impairment of BAK/BAX heterodimerization might prevent or delay the bleaching cascade in wild-type individuals. Vertebrate models show that BAK and/or BAX knockdown and knockout is a successful strategy to prevent PCD, even upon increased pro-dead signaling (*13*, *15*, *16*), and targeting BAK may be a more successful strategy than targeting BAX to avert PCD (*17–19*).

Many associated pathways have been proposed to play a role in coral bleaching and improved thermal tolerance (*5*, *6*, *8*, *20–22*). This complex physiology leads to a range of cellular and molecular consequences, including the accumulation of (oxidative) DNA damage in heat-stressed corals, which can be exacerbated by thermal preconditioning or by the addition of external antioxidants (*20*, *23*, *24*). The ROS-dependent bleaching theory postulates that during heat stress, symbionts are the main source of reactive oxygen species (ROS) which, unless scavenged by the host antioxidant defense, are detrimental to the host cell and trigger symbiosis disruption (reviewed in (*25*)). Increased DNA damage and the resultant compromised genome stability could be trade-offs to “forced” symbiosis stability in heat-stressed corals, such as those with manipulated gene expression.

In this work, we analyzed the connection between BAK manipulation and representative genes in antioxidant defense, DNA repair, programmed cell death, and AMPK and mTOR signaling pathways. We show that coral thermal resilience can be improved by the downregulation of a single gene, eliciting a complex molecular response that mitigates oxidative DNA damage in heat-stressed corals. While the precise mechanisms behind this improved phenotype remain unknown, we propose a novel pathway (AMPK/Foxo3a/Nrf2) that affects multiple pro-life and antioxidant genes involved in coral response to heat stress.

## Methods

### Coral collection and experimental setup

Twelve colonies of *Pocillopora acuta* (∼ 15 to 20 cm in diameter) were collected in 2022 and 2023 at 3 different sites in Kāneʻohe Bay, Hawaiʻi to maximize the likelihood of gathering distinct genotypes. Colonies were fragmented into ∼7 cm nubbins and maintained in shallow flow-through seawater tanks under natural light until the start of the experiment. The experimental setup is depicted in Figure S1.

### siRNA-mediated knockdown

The experiment was conducted as described in (*20*) with minor modifications. Freshly cut coral fragments (∼ 3 cm) were placed into 20 ml cultivation vials, completely submerged in a flow-through tank with ambient-temperature seawater, and left to acclimatize overnight. This step is crucial for the siRNA transfection to be successful. The next morning, vials were moved into a precise temperature-controlled illuminated water bath (280 series waterbath, Thermo Scientific with Hydra 64 HD LED Reef Light, AquaIllumination). Seawater was removed without disturbing the corals, and the siRNA transfection was carried out according to the manufacturerʻs instructions using a transfection mix consisting of 20μl of 10nM siRNA (siBAK targeting pa-BAK mRNA and siNTC designed as a negative control with no known target in the *P. acuta* mRNA sequence database, see Table S1 for sequences) and 16μl INTERFERin® reagent (Polyplus) in 400μl of 0.2μm filtered seawater pipetted directly onto the exposed coral. After 2 minutes, the coral fragment was covered in 5ml of 0.2μm filtered seawater and incubated at ambient temperature for 6 hours. 30 minutes after the transfection, soft aquarium airline PVC tubing (4 mm in inside diameter, NACX) was inserted into each cultivation vial, and the seawater was aerated with approximately 1 bubble every 3 seconds to prevent hypoxic stress without disturbing the corals. After the 6-hour incubation, coral fragments were removed from the transfection mixture, placed in a 6-well plate and completely submerged in a flow-through seawater tank. 24 hours after the beginning of the siRNA transfection, corals were exposed to acute heat stress ramping from ambient temperature (fluctuating between 25.3 to 26.5, depending on the day of the experiment) to 32°C over 2.5 hours before being held constant at 32°C. After 48h in the stress treatment, we evaluated fragments with microscopy (see below), and small subsamples were flash-frozen in liquid nitrogen and maintained at –80°C until extractions for gene expression and DNA damage analyses. We re-evaluated the corals using microscopy at 96h in the stress treatment. The experiment was run across 6 trials with up to 3 genotypes at each time, totaling 16 different individual siRNA-delivery attempts. Four corals were used twice for the experiment (with the first transfection not being successful) in order to achieve successful downregulation in 10 different genotypes.

### Bleaching rate assessment

We evaluated bleaching using confocal microscopy analysis (Zeiss LSM-710) following (*5*). The natural fluorescence of algal-derived chlorophyll a was normalized to the natural fluorescence signal of animal-derived GFP protein and used as a proxy for the symbiont density in the coral tissue (*5*, *26*). Coral fragments were measured the day before siRNA transfection and 48 and 96 hours post-heat stress. Three snaps of the top side of each coral were taken which covered the entire fragment area. Ten circular regions of interest (ROI) were analyzed on the focal plane of each picture (Fiji, (*27*)), avoiding the polyp mouth and tentacles that emit intense GFP-like signal. All ten ROIs were averaged and one value per snap was used for the statistical analysis. Three snaps were analyzed for each coral per timepoint per treatment.

### DNA damage assay

DNA was extracted from corals at the 48h timepoint (72h after the transfection) using Quick-DNA/RNA^TM^ Microprep Plus Kit (Zymo Research). The level of 8-Hydroxy-2’-deoxyguanosine (8-OhdG) – a proxy for oxidative DNA damage – was analyzed in corals using DNA Damage Competitive ELISA Kit (Invitrogen) and MyCurveFit.com software as described in (*20*).

### Gene expression analysis

Gene expression was analyzed as described in (*5*) with minor changes. RNA was extracted from samples collected at the 48h timepoint (72h after the transfection) using Quick-DNA/RNA^TM^ Microprep Plus Kit (Zymo Research) including the DNAse I step. 1000 ng of DNA was transcribed with a High-Capacity cDNA reverse transcription kit (Thermo Fisher Scientific) with the addition of 20 U of Murine Rnase Inhibitor (New England Biolabs). qPCR reactions were run in 12 μl with PowerUp^TM^ SYBR^TM^ Green Master Mix (Applied Biosystems) for 40 cycles according to the manufacturerʻs instructions. The efficiency of each primer pair was tested with a dilution curve followed by a melting curve analysis. Pa-EF-1 (elongation factor 1a) was used as a normalizing gene based on the analysis published in (*5*). Expression of target genes in siBAK-treated corals was calculated relative to the siNTC-treated corals with the ΔΔCt method.

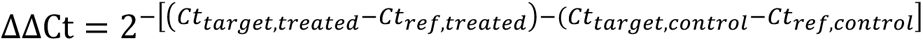

Each analysis included one sample per treatment for each colony.

For the siBAK treatment response, only samples from corals with a successful pa-BAK knockdown were analyzed, highlighting the effect of the pa-*BAK* downregulation on the downstream gene expression. For the correlation analysis, the data from all 16 colonies (including those where the siBAK treatment was not successful) were used to describe the global gene profiles linked to pa-BAK and to improve statistical power. We then re-evaluated the correlation in the subset of corals with successful siBAK knockdown to emphasize genes that respond directly.

We analyzed the expression of the following genes: pa-GR (Glutathione reductase), pa-OGG1 (8-oxoguanine DNA glycosylase), pa-FOXO3a (forkhead box O3), pa-HO-1 (heme oxygenase), pa-BI-1 (Bax inhibitor 1), pa-Bcl-2 (B-cell lymphoma 2), pa-BAK (Bcl-2 homologous antagonist/killer) pa-BAX (Bcl-2 associated X protein), pa-Cas3 (Caspase 3), pa-GABARAP (GABA type A receptor-associated protein), pa-LC3 (microtubule-associated proteins 1A/1B light chain 3B), pa-AMPK (AMP-activated protein kinase), and pa-mTOR (mechanistic target of rapamycin). Sequences of all primers used in this study are listed in Table S1.

### Statistical analyses

All statistical analyses and visualizations were conducted in R, Version 2023.06.1+524 (*28*). We evaluated normality and heteroscedasticity for each dataset. The efficiency of siBAK knockdown was analyzed with a one-sample t-test, comparing exprsesion to 1. The Chl a/GFP ratio calculated from confocal microscopy was analyzed using a generalized linear mixed model in *lme4* (*29*) with treatment and time as main effects and colony as a random effect (ratio∼treatment*time+(1|coral).

Bleaching rate was calculated for each treatment as the slope of the fit between time (hours) and Chl A/GFP and used to calculate the tolerance gain (the change in bleaching after siBAK knockdown) using the following formula:

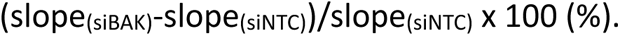

pa-BAK knockdown efficiency was calculated as the gene expression in siBAK normalized to siNTC via the ΔΔCt method, retracted from one, and expressed in percentages using the following formula:

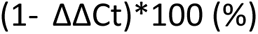

We calculated the tolerance gain for each coral and compared it to the pa-BAK knockdown efficiency using a Pearson correlation.

The oxidative DNA damage marker 8-OhdG was compared using the Paired Studentʻs T-test. We used hierarchical clustering using the ‘complete’ method in the R package *hclust* to find relative relationships between genes. The relationship between pa-BAK expression at 48 hours and other genes was evaluated using a Pearson correlation. This correlation was examined in all siBAK-treated corals (n = 16) and separately only in the genotypes with efficient BAK knockdown (n = 10). A one-sample t-test was used to evaluate if the expression of each gene was different from 1 in genotypes with efficient BAK knockdown.

## Results

### pa-BAK knockdown improves bleaching resilience of corals

We conducted the siRNA delivery in 16 coral fragments during 6 separate experiments and downregulated the expression of the pa-*BAK* gene in 10 different colonies (55.6%). We analyzed pa-*BAK* expression at 48 hours post-heat stress and designated each trial a successful knockdown when the pa-*BAK* expression was less than 80% of siNTC-treated corals. This timepoint was a total of 72 hours post-siRNA delivery (including the 24 hours of transfection and recovery), which is generally considered the latest timepoint for mRNA level analysis because the induced mRNA downregulation loses efficiency over time (Interferrin™ protocol, Polyplus). We chose this timepoint as the compromise between the time we can still observe efficient downregulation and the onset of the phenotypic shift in the bleaching rate. Expression after knockdown averaged 59% ± 24% (mean ± SD, range: 2% to 79%, Figure 1A) of control.

**Figure 1.**
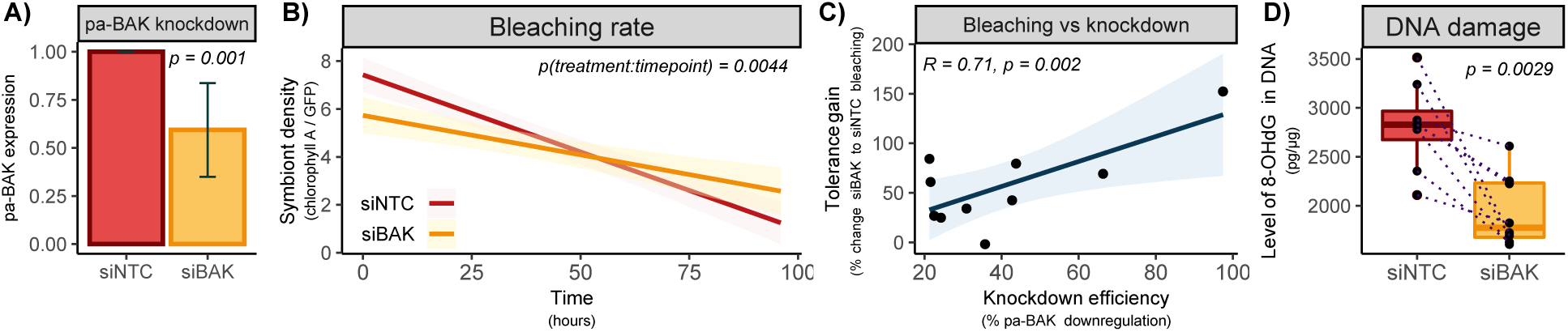
pa-BAK knockdown improves thermal resilience and decreases oxidative damage in heat-stressed corals. A) Successful siRNA-mediated downregulation of pa-BAK in *P. acuta* corals. The expression is normalized to the control treatment (siNTC) by the ΔΔCT method. The bar represents standard deviation, n =10. B) Bleaching rate in heat-stressed corals with successful pa-BAK knockdown (siBAK) compared to their controls (siNTC), n = 10. C) Correlation between the tolerance gain (calculated as the percentual difference between bleaching rates of treated (siBAK) and control corals (siNTC)) and the magnitude of pa-BAK knockdown (calculated as percentual downregulation compared to the control), R stands for Pearson correlation coefficient, n = 10. D) The level of oxidative DNA damage marker (8-OHdG) in DNA of heat-stressed corals with successful pa-BAK knockdown (siBAK) and controls (siNTC), n = 10.

We then analyzed the bleaching rate in heat-stressed corals by comparing Chla (symbiont-derived signal) to GFP (host-derived signal) ratio over time following (*5*, *26*). Corals were scanned the day before the siRNA delivery, 48 hours into the acute heat stress (i.e., 72 hours post-siRNA delivery), and 96 hours into the acute heat stress (i.e., 120 hours post-siRNA delivery). Exposure to heat stress significantly decreased Chla:GFP ratio (p_(time)_ < 0.001, Figure 1B and S2A) and corals with downregulated expression of pa-*BAK* bleached 51% slower than controls (p_(treatment:timepoint)_ = 0.004). Corals in which BAK downregulation was not successful did not show any shift in bleaching rate (Figure S2B).

The degree of improvement in bleaching tolerance – the tolerance gain – was positively correlated with the magnitude of pa-*BAK* downregulation (R = 0.77, p = 0.009, Figure 1C); corals with more pronounced pa-BAK knockdown bleached at a slower rate (normalized to their individual controls) during acute heat stress. This relationship was driven by the siRNA-mediated knockdown; the tolerance gain did not correlate with pa-BAK expression in heat-stressed corals in which BAK was not successfully downregulated (R = 0.3, p = 0.56, Figure S2C).

### Markers of oxidative DNA damage decrease in corals after paBAK knockdown

Corals with effective pa-BAK knockdown showed significantly decreased levels of oxidative DNA damage (8-OhdG, 8-hydroxy-2ʻ-deoxyguanosine) after 48 hours in heat stress, 72 hours post-siRNA delivery (paired Studentʻs T-test, p = 0.0029, Figure 2D) and all genotypes tested showed less DNA damage in the siRNA treatment. However, the level of oxidative DNA damage did not correlate with the tolerance gain (R = –0.59, p = 0.12, Figure S2D) or with the extent of the pa-BAK downregulation (R = 0.36, p = 0.39, Figure S2E)

**Figure 2.**
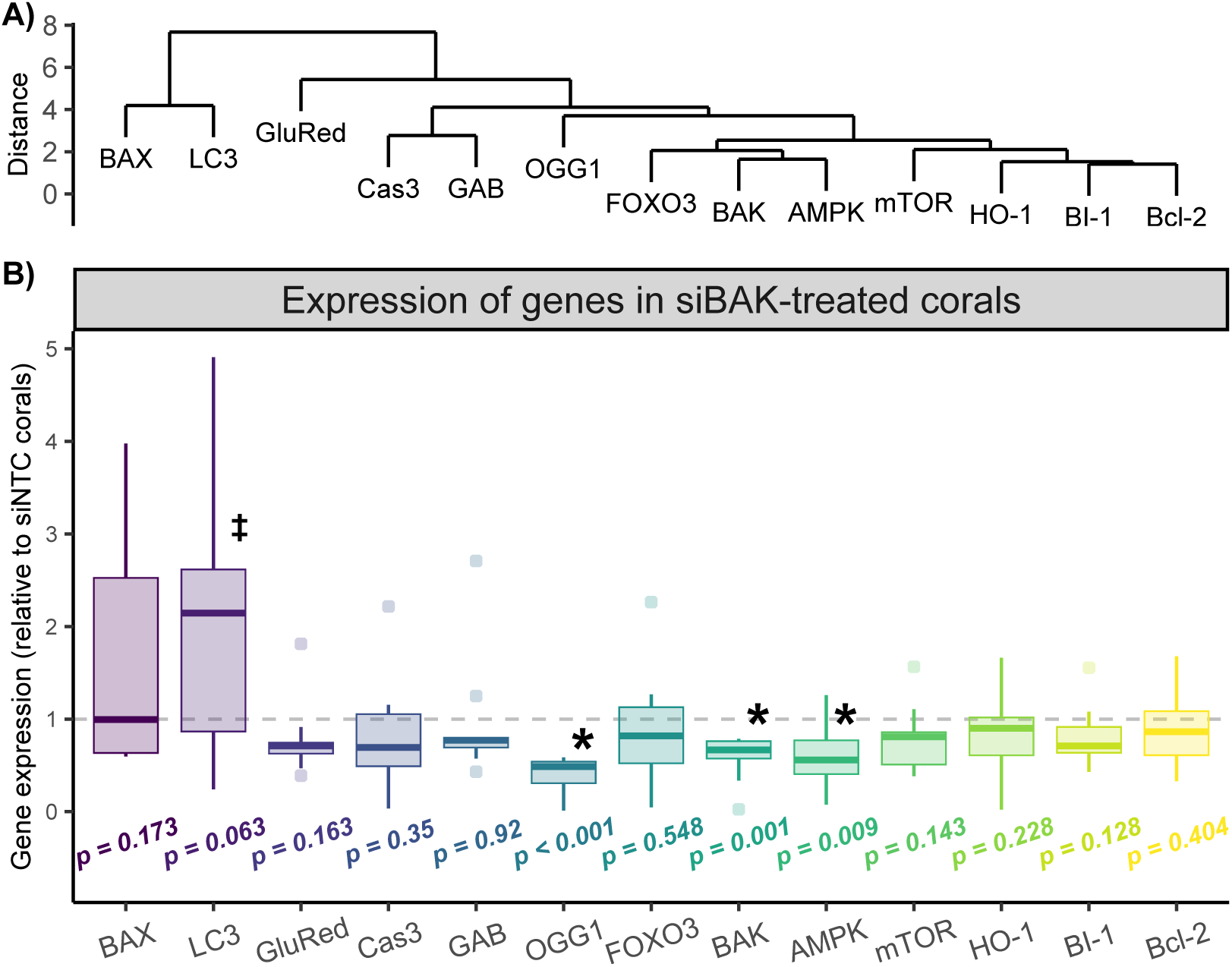
Expression of genes in heat-stressed *P. acuta* corals. A) Hierarchical clustering based on the gene expression in all heat-stressed corals shows BAX and LC3 having relatively discrete profiles, while the rest of the genes respond to heat stress more closely. n = 16. B) Gene expression in corals with downregulated pa-BAK. * represents p-values lower than 0.05, ‡ represents p-values between 0.05 and 0.1, n = 10.

### Genes connected to programmed cell death and oxidative stress response correlated in heat-stressed corals

To better understand what cellular and molecular pathways are affected during the phenotypic change associated with BAK downregulation and heat stress in general, we examined the pairwise relationships of the expression of a variety of genes. Hierarchical clustering suggested that BAX and LC3 had relatively discrete expression profiles, while the rest of the genes were in a single node of the dendrogram, corresponding more closely (Figure 2A).

BAK and BAX can compensate for each other under certain conditions (*17*, *30*). To eliminate this possibility, we examined the expression of pa-*BAX* in siBAK-treated corals (Figure 2B). While slightly upregulated, its expression was not different in pa-*BAK* downregulated corals (p = 0.173), and the expression of pa-*BAX* does not correlate with pa-*BAK* (R = – 0.11, p = 0.724, Figure 3). pa-*OGG1* and pa-*AMPK* expression were significantly downregulated in pa-*BAK* downregulated corals (p < 0.001 and p = 0.009, respectively), and pa-*LC3* was slightly upregulated (p = 0.063)(Figure 2B).

**Figure 3.**
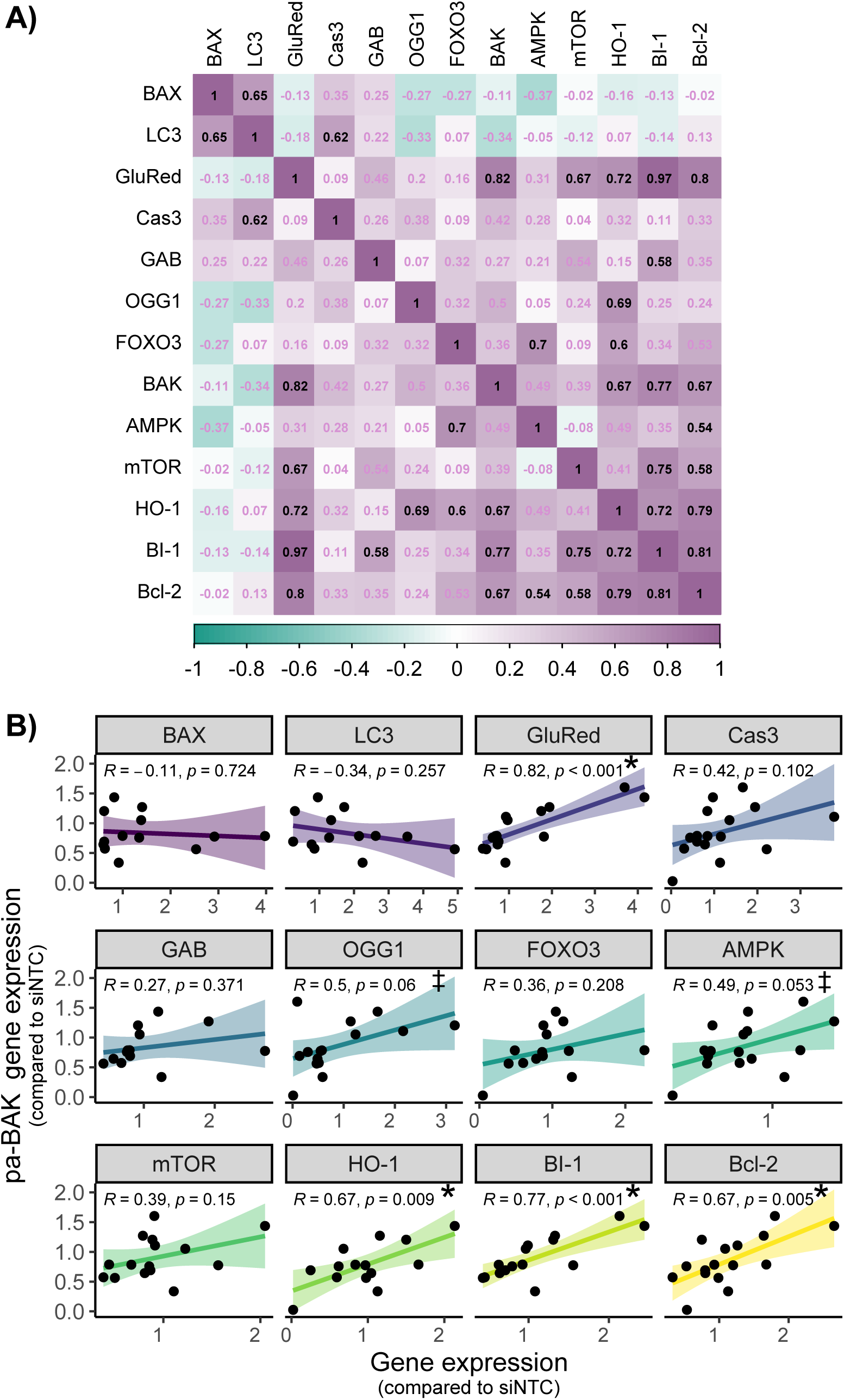
Gene expression correlations in all heat-stressed *P. acuta*. A) Heat map of gene expression correlations. Numbers represent Pearson correlation values, significant (p<0.05) in black, and non-significant (p>0.05) in pink, n = 16. B) Relationship between pa-BAK gene expression and expression of other genes. Expression is normalized to the control treatment (siNTC) by the ΔΔCT method. P stands for Pearson correlation coefficient, * represents p-values lower than 0.05, ‡ represents p-values between 0.05 and 0.1, n = 16.

Pairwise correlation of all genes (Figure 3A, Table S2, Figure S3) showed a group of genes (pa-*GR*, pa-*BI-1*, pa-*Bcl-2*, and pa-*HO-1*) strongly correlate with each other and with pa-*BAK* (Pearson R >0.67, p <0.009). Hierarchical clustering identified three of these genes (pa-*BI-1*, pa-*Bcl-2*, and pa-*HO-1*) as having the most similar expression profiles. Pa-*OGG1* and pa-*AMPK* almost significantly correlated with the pa-*BAK* expression (R = 0.5, p = 0.06 and R = 0.49, p = 0.053, respectively). Pa-*BI-1* strongly correlated with pa-*mTOR* (R = 0.75, p = 0.001) and pa-*foxo3* was strongly correlated with pa-*AMPK* (R = 0.7, p = 0.005).

Multiple pairs of genes showed moderate correlation. Pa-*OGG1* and pa-*HO-1* (R = 0.69, p = 0.009), pa-*mTOR* and pa-*GR* (R = 0.67, p = 0.006), pa-*LC3* and pa-*BAX* (R = 0.65, p = 0.017), pa-*LC3* and pa-*Cas3* (R = 0.62, p = 0.025), pa-*HO-1* and pa-*foxo3* (R = 0.6, p = 0.023), pa-*mTOR* and pa-*Bcl-2* (R = 0.58, p = 0.023), pa-*GABARAP* and pa-*BI-1* (R = 0.58, p = 0.037), and pa-*AMPK* and pa-*Bcl-2* (R = 0.54, p = 0.031).

When we analyzed the correlation of each gene with pa-BAK using only corals that were successfully downregulated after the siBAK treatment, there were no significant relationships (data not shown).

Pa-*OGG1* and pa-*foxo3* gene expression were positively correlated with the level of oxidative DNA damage marker in heat-stressed corals (R = 0.64, p = 0.088 and R = 0.67, p = 0.069, respectively, Figure S4A). pa-*AMPK* and pa*-OGG1* expression mildly negatively correlated with tolerance gain (R = –0.52, p = 0.041, and R = – 0.48, p = 0.068, respectively, Figure S4B).

## Discussion

Coral thermal tolerance is a complex trait driven by the interaction between cnidarian host and photosynthetic symbionts, making it especially difficult to describe and manipulate. As coral reefs face an increasingly stressful future of warming oceans driven by anthropogenic change, developing tools and a basic understanding of these mechanisms is critical for conservation and management. Here we show that the manipulation of a single gene slows coral bleaching, decreases oxidative DNA damage, and creates an extensive cellular cascade with potentially transformative effects.

Programmed cell death pathways are one of the common mechanisms of coral-algal symbiosis disruption (*4–6*, *8*, *10*, *11*). There is a well-documented cross-talk between apoptosis and autophagy, and several proteins such as Bcl-2, BAK, or BAX are described to have the power to activate/prevent both pathways, depending on particular cellular conditions (*12*, *31*). The expression ratio of pro-life Bcl-2 to pro-death BAK and BAX increases during acute heat stress in *P. acuta* and *Acropora millepora* and is significantly higher in corals with higher thermal resilience (*5*, *6*). Inhibition of pa-Bcl2 leads to the loss of the beneficial phenotype (*5*). Since the disruption of symbiosis in corals is the major response mechanism to various stress conditions (*32*), it is more meaningful – and also more challenging – to search for gene manipulations that can enhance, not disrupt the stability of coral-algal symbiosis under thermal stress. Experiments in vertebrates have shown that BAK or/and BAX downregulation is a successful strategy to prevent programmed cell death pathways (PCD) in cells (*13*, *15*, *17*) and our results demonstrate that pa-BAK knockdown is sufficient to slow down but not fully prevent heat-induced coral bleaching (Figure 1B). This may be due to the incomplete nature of expression decline using si-RNA mediated knockdown, leaving enough pa-BAK to heterodimerize with pa-BAX and activate the PCD, or that other but less powerful mechanisms compensate for the loss of the pro-death PCD signaling. It is also possible that symbionts actively leave coral tissue during acute heat stress, independent of the host cell mechanisms.

Heterodimerization of BAK and BAX and the subsequent permeabilization of the mitochondrial membrane is considered a non-reversible point in the apoptotic cascade (*13*): it triggers a self-destroying cellular mechanism based on the expression and activation of multiple executioner proteases of the family of caspases (*12*). The detailed mechanism of BAK/BAX’s role in the activation of autophagy is not yet well understood, but it is also presumed that they are the main trigger before the execution of the pathway (*12*, *13*, *31*). Several studies showed that depletion of both proteins is necessary to efficiently block the execution of the programmed cell death pathway (*16*, *30*), potentially because BAK can be upregulated in BAX-depleted cells, efficiently compensating for loss of function (*17*). In our experiment, we found an improvement in symbiosis maintenance under heat stress but did not observe a compensatory relationship between BAX and BAK upon knockdown, suggesting that BAK has a unique, BAX-independent role in the disruption of the coral host-algal symbiosis. This pattern is supported by the observation that the more effective the BAK knockdown was, the slower the coral bleached during heat stress (Figure 1C). Since this correlation is absent in corals with unsuccessful pa-*BAK* knockdown (Figure S2B), we conclude that the level of pa-*BAK* expression is not an isolated marker of symbiosis stability, but its inhibition improves this phenotype. BAK’s distinctive BAX-independent role has also been described in model organisms (*18*, *19*).

Coral thermal tolerance and stress response are intricate traits that employ myriad cell pathways and affect coral physiology and metabolism (*5*, *20–22*, *33–35*). The complexity of these processes means there are likely to be off-target consequences of ‘forced’ thermal tolerance, so the evaluation of other, potentially detrimental traits, is critical from a basic and applied perspective. Thermal stress causes the accumulation of oxidative DNA damage markers in coral tissue which can be prevented through preconditioning (*20*). We hypothesized that heat-stress-induced DNA damage could trigger PCD and lead to subsequent bleaching (*5*, *20*). If the accumulated oxidative DNA damage in coral cells is the result of non-functional symbiosis and/or if it signals for the activation of PCD, we expect to see similar or even increased DNA damage marker levels in pa-*BAK* downregulated corals compared to the controls as the damage would be accumulating in cells otherwise programmed to self-destroy. Surprisingly, we observed a significantly decreased level of oxidative DNA damage marker in BAK knockdown corals undergoing 48 hours of acute heat stress (Figure 1D). This oxidative mutation is usually recognized and repaired by DNA glycosylases, most commonly OGG1 (8-oxoguanine DNA glycosylase), via the process of base excision repair (BER) (*36*). In alignment with the observed decrease in oxidative DNA damage, the expression of pa-*OGG1* also significantly decreased in pa-BAK downregulated heat-stressed corals (Figure 2B), and the level of pa-*OGG1* expression positively correlated with the DNA damage extent and bleaching rate (FigureS4). The slower corals bleached, the less pa-OGG1 was expressed, and the lower DNA damage was detected (in a correlative, not causal relationship). In addition, there was a weak positive correlation between pa-*OGG1* and pa-*BAK* expression (R = 0.5, p = 0.06, Figure 3A, Table S2). If the si-BAK treatment led to the improvement of oxidative DNA damage repair mechanisms, we would expect to observe increased levels of pa-*OGG1* in pa-BAK downregulated coral cells, but the results show the opposite. Decreased expression of pa-*OGG1* in corals with improved symbiosis stability under heat stress suggests that oxidative DNA damage is prevented, not repaired in pa-*BAK* knockdown corals.

Neither of the antioxidant genes, Glutathione reductase (GR) and Heme oxygenase 1 (HO-1), were expressed differently in BAK-downregulated corals (Figure 2B). However, pa-*HO-1* showed a moderate correlation with pa-*OGG1* gene expression (Figure 3A), which was very strong when only successfully BAK-downregulated corals were considered (R = 0.91, p < 0.001, data not shown). Heme oxygenase 1 is an inducible form of heme oxygenase enzyme that catalyzes heme breakdown. Its byproducts possess anti-inflammatory, anti-apoptotic, and antioxidant properties and a truncated form of HO-1 with low enzymatic activity acts as a transcriptional factor in the nucleus during stress response (reviewed in (*37*)). OGG1 can also induce gene transcription, although the mechanisms remain largely unclear (*36*). It is thus possible that one protein regulates the expression of the other gene, or that they are both controlled by a third party. HO-1 knockdown leads to downregulation of OGG1 in irradiated mice lungs (*38*); however, studies also show that both HO-1 and OGG1 are positively and directly regulated by the Nrf2 factor (*39*, *40*).

We previously hypothesized that more thermally resilient corals enhance their antioxidant and DNA damage defense systems via the Nrf2/ARE pathway (20). BI-1 mediated upregulation of glutathione reductase improves oxidative DNA damage defense in preconditioned (thus thermally more resilient) P. acuta. Similarly, we also found a strikingly high correlation between pa-BI-1 and pa-GR expressions in heat stress (Figure 3A and S3, Table S2) confirming this previously established relationship. pa-Bcl-2, a major pro-life gene defined in several programmed cell death pathways, is also highly correlated with the pa-BI-1 and pa-GR expression profiles, all of which are correlated with pa-HO-1. Bcl-2 expression can be controlled by the Nrf2 pathway (41) but overexpression of Heme oxygenase 1 is enough to induce the expression of Bcl-2 in a rat heart, suggesting HO-1 might be an intermediary in the regulation (42). Additionally, HO-1 can regulate the expression of Nrf2 in what appears to be a positive feedback loop (37) which points to a complex regulatory network surrounding the Nrf2 stress-response pathway. Nrf2 is a pleiotropic factor, and it may thus play a major role in acclimatization to environmental stress in corals with improved symbiosis stability impacting multiple important defense mechanisms in the organism. It would be interesting to further study the involvement of this pathway in coral thermal resilience.

Foxo3a is linked to the modulation of stress responses upon oxidative stress, DNA damage, or nutrient shortage (reviewed in (*43*)). Its activity is primarily controlled by post-translational modifications, which determine subcellular localization. We found the expression of pa-*foxo3* positively correlated with pa-*AMPK* and pa-*HO-1* (Figure 3A, and S3, Table S2). AMPK activates Foxo3a mostly as a response to nutrient stress which leads to transcription of autophagy-related genes (*43*). Lower expression of pa-*AMPK* could lead to decreased pa-*foxo3* expression in BAK-knockdown corals, which would lead to lower expression of autophagy/symbiophagy-related genes. We previously identified symbiophagy rather than apoptosis as the probable pathway involved in thermal resilience in *P. acuta* (*5*). AMPK has been also proposed to boost the Nrf2-mediated HO-1 expression (*44*), and Foxo3a putatively regulates the keap1/Nrf2 pathway, likely activating or repressing it based on specific conditions (*45*). Given the results of this study, we propose that the main pro-life and antioxidant genes (pa-*GR*, pa-*BI-1*, pa-*Bcl-2*, and pa-*HO-1*) are regulated through the AMPK/Foxo3a/Nrf2 signaling. However, AMPK can be (probably indirectly) activated by mitochondria-derived ROS (*46*), thus we cannot entirely exclude that the AMPK/Foxo3a regulation is independent of the putative Nrf2-mediated control of pa-GR, pa-BI-1, pa-Bcl-2, and pa-HO-1. Therefore, their correlation would not be causal and would exist in parallel and the two major signaling pathways – Nrf2 and AMPK would act independently in heat-stressed corals. This hypothetical ambiguity needs to be further examined using the tools of functional genetics to overcome the limitations of describing fine-tuned signaling of cellular response pathways that are primarily regulated via posttranslational modifications rather than gene expression.

This hypothesized signaling network still does not explain the mechanism causing improved thermal resilience and oxidative damage defense in pa-BAK downregulated corals. Recent research shows that many proteins of the Bcl family (including BAK and BAX) have different non-apoptotic functions, often related to mitochondrial structure and functioning (*47*). In normal conditions, BAK – unlike BAX – is localized inside of mitochondria with its C-terminal region inserted in the lipid bilayer (*48*). The observed phenotypical advantage of pa-BAK downregulation changes in gene expression could be due to the changes in the pool of “available” single BAK protein inserted in the mitochondrial membrane. The presence of BAK is necessary for the formation of the so-called mitochondrial permeability transition pores (mPTP) (*49*). Their regular openings help maintain healthy mitochondria homeostasis as they control the release of excessive ROS to the cytoplasm where it affects stress signaling and oxidative damage response pathways (*49*, *50*).

In some cases, released ROS triggers the formation of mPTP and subsequential ROS release from neighboring mitochondria, activating a positive feedback loop called ROS-induced ROS release (RIRR), which plays a strategic role in extensive signaling and may lead to oxidative cellular damage and cell death via a PCD pathway (*49*). The loss of BAK and BAX causes changes in the mPTP permeability, prevents mitochondrial swelling and the cells resist cell death (*50*). The inner membrane still undergoes stress-related reorganization, so the possible long-term outcomes of such forced survival on mitochondrial and cellular health are important areas of future research. Besides energy production, mitochondria play a vital role in intra– and extracellular communication, and their impaired function is connected to a multitude of symptoms and degenerative diseases (reviewed in (*51*)). These additional questions must be resolved to determine if the siBAK phenotype can be truly deemed beneficial for corals in the long term.

Algal symbionts co-exist within the host cell in symbiosomes, which are believed to be late lysosomes in the state of arrested phagocytosis (*52*). LC3 is often analyzed as a proxy for the amount and/or size of autophagosomes in the cell, as it is incorporated into the membrane of newly emerging phagosomes during the initialization phase (*53*). Here, pa-*LC3* was slightly increased in BAK-knockdown corals which might point to the increased symbiont stability or possibly to an increased turnover of symbionts during acute heat stress (increased level of LC3 can be interpreted as an increased number, size, or turnover of phagosomes and should not be interpreted in a definite manner by itself without additional experiments (*53*)). pa-*GABARAP*, another marker of phagosomes, did not respond to the treatment, and its expression was weakly correlated with pa-*BI-1*, suggesting it may be coincidental.

This work demonstrates that corals can become more resilient during acute heat stress via targeted manipulation of a single gene. It also shows that corals modulate various pathways as a response to single gene manipulation, probably to cope with possible harmful trade-offs of a prolongated symbiosis in heat stress. However, the correlative gene network described here relies on the inclusion of all corals in the analysis; most of the correlations are no longer significant when only corals with downregulated pa-*BAK* are considered. This suggests that we are observing and describing pathways generally involved in coral response to heat stress and not pathways that lead to improved symbiosis stability and DNA damage prevention in corals with impaired pa-*BAK* function, in particular. Our analysis also focused on a single timepoint, which means that the absence of correlation could be caused by the arbitrary choice of timepoint and thus we are missing the dynamic nature of stress response and signaling pathways in heat-stressed corals.

These data shed light on the network of regulatory pathways involved in coral response to acute heat stress. Our results are also are crucial for assisted evolution approaches to conservation and restoration that target more resilient individuals and uses them to support the health of coral reefs facing anthropogenic climate change (*54*, *55*).

## Acknowledgments

We acknowledge the profound connection between Native Hawaiian culture and coral reefs (*56*), including those found in Kāneʻohe Bay, Oʻahu, Hawaiʻi, where this research was conducted. We hope to honor the unique relationship between local communities and place, and the practice of mālama ʻāina (caring for and protecting the land).

EM conceived the experiments, conducted research, analyzed and visualized data, and wrote the manuscript. CS conducted research, visualized data, and wrote the manuscript. CD analyzed data and wrote the manuscript. This work was funded by the National Science Foundation (IOS-2041401). Corals were collected under permit SAP 2023-31. The authors declare they have no competing interests. All data needed to evaluate the conclusions in the paper are present in the paper and/or the Supplementary Material. We would like to thank Kira Hughes, Khalil Smith, and Joshua Hancock for assistance with coral husbandry, experimental setup, and project administration and the Coral Resilience Lab for helping with coral collection and maintenance.

## Supplemental material

**Figure S1.**
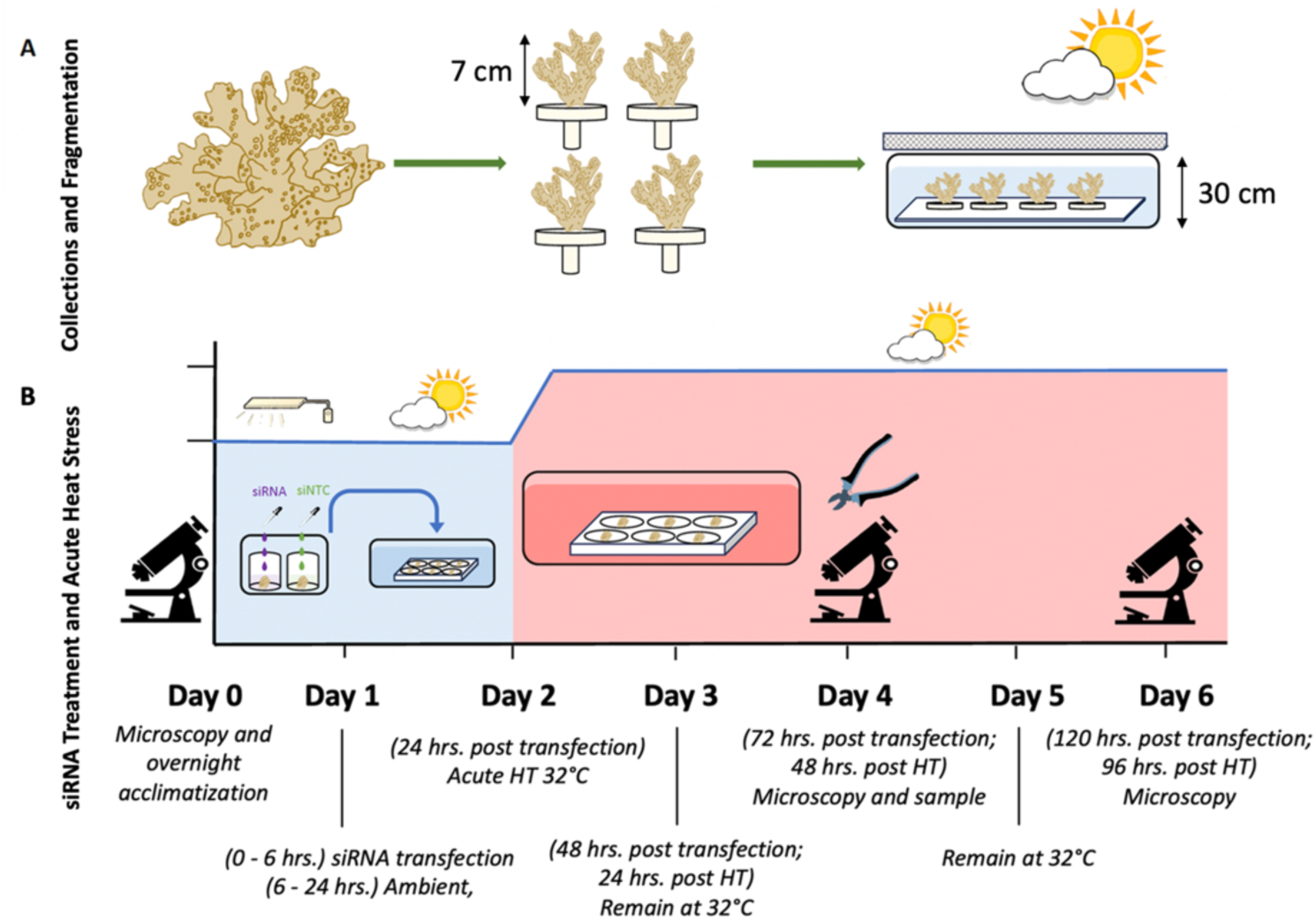
Experimental setup layout.

**Figure S2.**
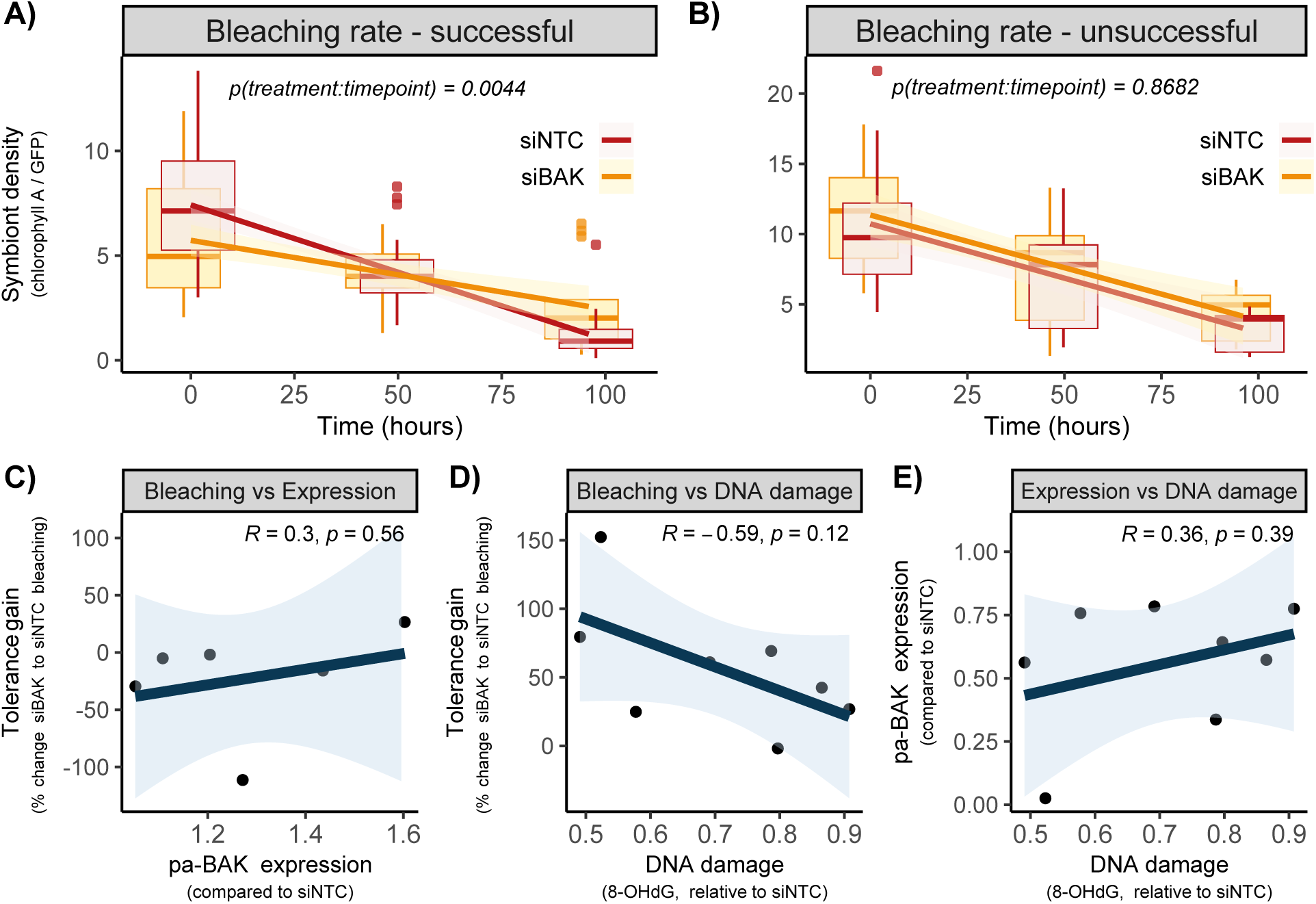
Physiological and cellular response to heat stress in *P. acuta* corals. A,B) Bleaching rate in heat-stressed corals with successful (left) and unsuccessful (right) pa-BAK knockdown compared to their controls (siNTC). The boxplots show the median of the data, first and third quartile, and outlying datapoints. n = 10 and 6, respectively. C) Correlation between tolerance gain and pa-BAK expression in corals with unsuccessful pa-BAK knockdown. D, E) Correlation between DNA damage and tolerance gain (D) or pa-BAK expression (E) in corals with successful pa-BAK knockdown. R stands for Pearson correlation coefficient.

**Figure S3.**
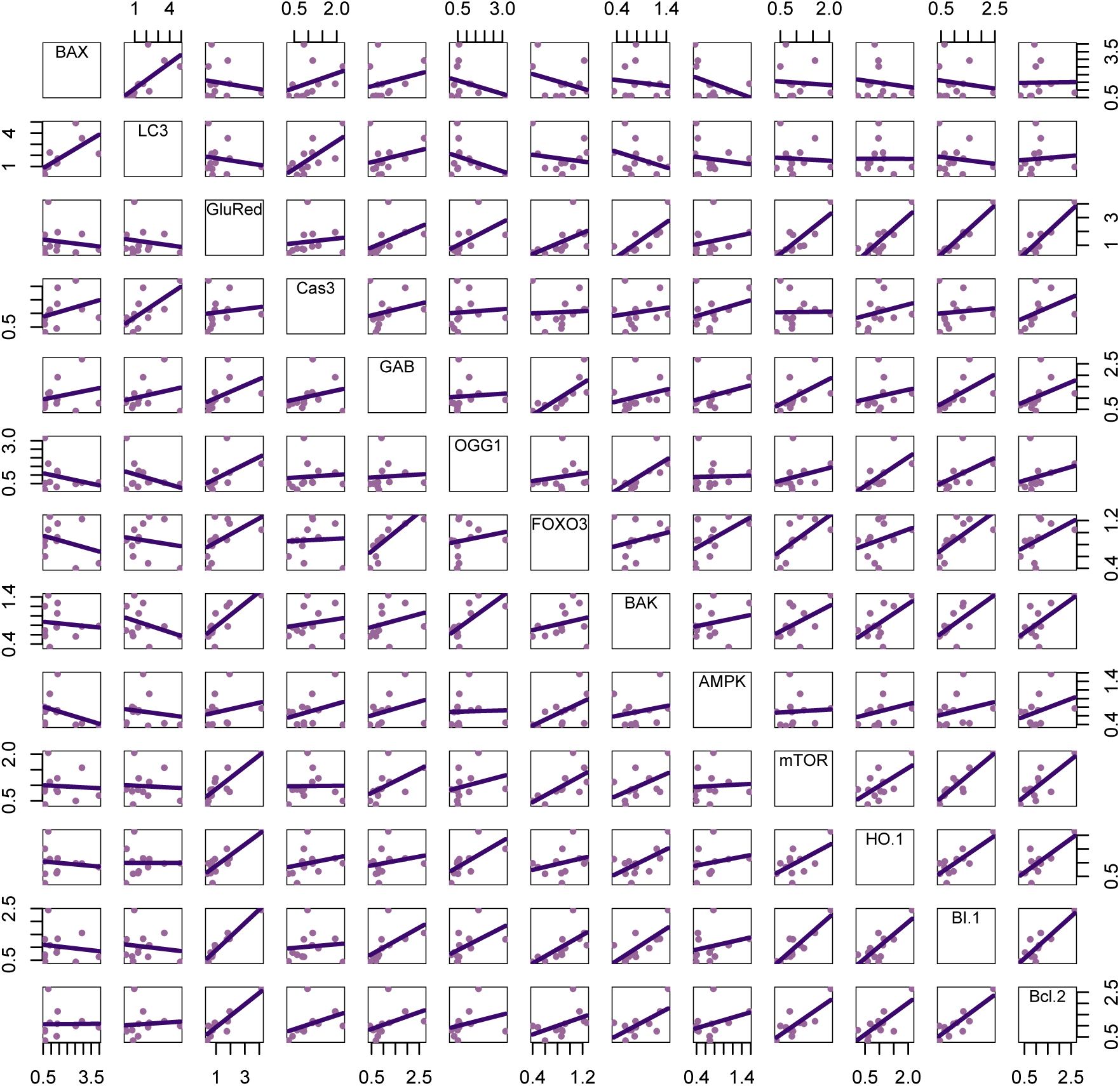
Scatterplot matrix for pairwise gene expression correlations. The expressions are normalized to the control treatment (siNTC) by the ΔΔCT method, n = 16.

**Figure S4.**
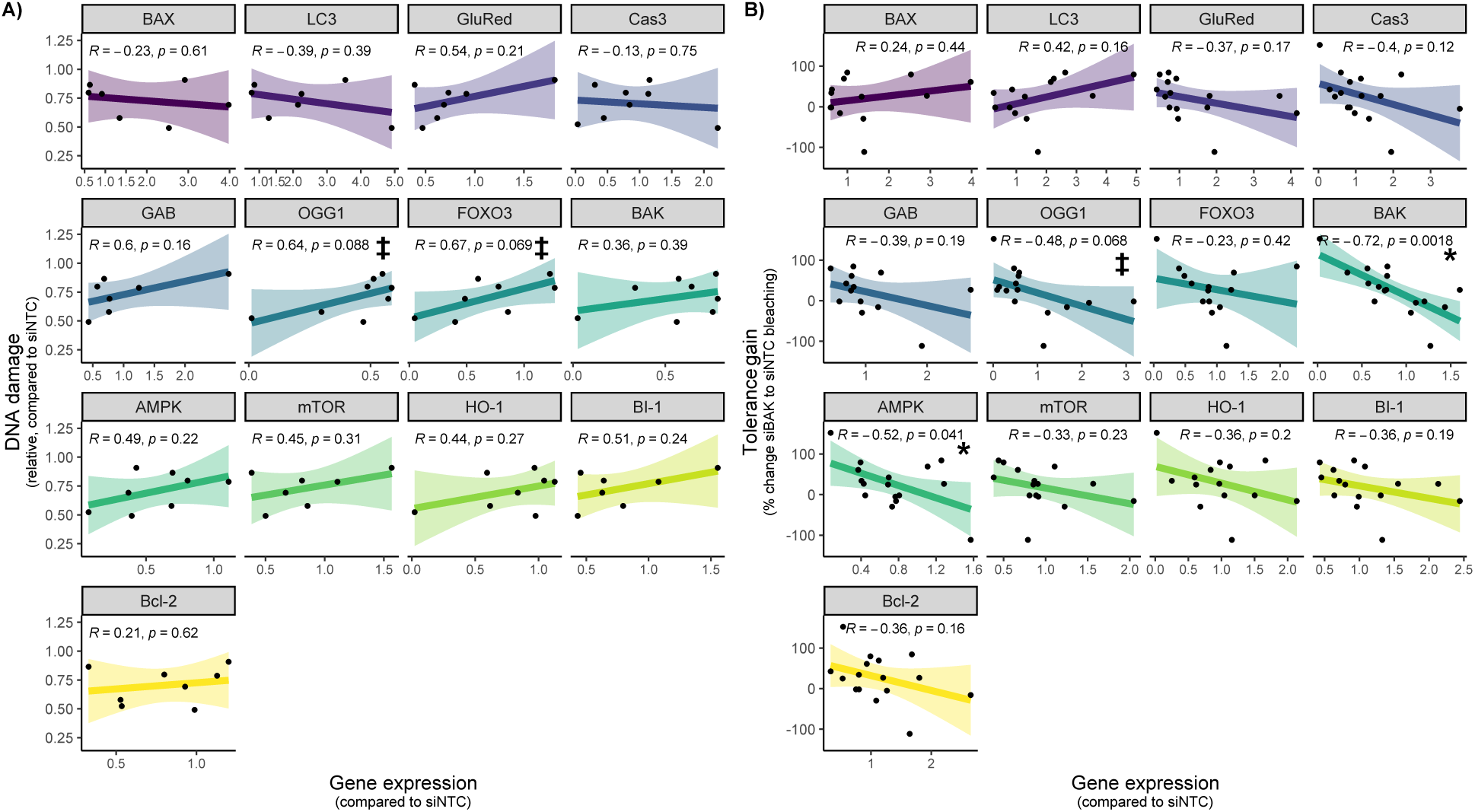
Relationship between physiological and molecular traits in heat-stressed *P. acuta* corals. Correlation between gene expression and DNA damage level (A) or tolerance gain (B), respectively. The expressions are normalized to the control treatment (siNTC) by the ΔΔCT method. P stands for Pearson correlation coefficient, * represents p-values lower than 0.05, ‡ represents p-values between 0.05 and 0.1, n = 16.

